# sc-REnF:An entropy guided robust feature selection for clustering of single-cell rna-seq data

**DOI:** 10.1101/2020.10.10.334573

**Authors:** Snehalika Lall, Abhik Ghosh, Sumanta Ray, Sanghamitra Bandyopadhyay

## Abstract

Many single-cell typing methods require pure clustering of cells, which is susceptible towards the technical noise, and heavily dependent on high quality informative genes selected in the preliminary steps of downstream analysis. Techniques for gene selection in single-cell RNA sequencing (scRNA-seq) data are seemingly simple which casts problems with respect to the resolution of (sub-)types detection, marker selection and ultimately impacts towards cell annotation. We introduce **sc-REnF**, a novel and **r**obust **en**tropy based **f**eature (gene) selection method, which leverages the landmark advantage of ‘Renyi’ and ‘Tsallis’ entropy achieved in their original application, in single cell clustering. Thereby, gene selection is robust and less sensitive towards the technical noise present in the data, producing a pure clustering of cells, beyond classifying independent and unknown sample with utmost accuracy. The corresponding software is available at: https://github.com/Snehalikalall/sc-REnF

## Introduction

In recent times technological advances have made it possible to study RNA-seq data at single cell resolution^1^. Single cell RNA sequencing (scRNA-seq) is a powerful tool to capture gene expression snapshots in individual cells. Cell type detection is one of the fundamental steps in downstream analysis of scRNA-seq data^2^. A widely used approach for this is to cluster the cells into different groups, and determine the identity of cells within the individual groups/clusters^3, 4^. This provides an unsupervised method of annotating the different cell types present in the large population of scRNA-seq data^5–8^.

Starting from raw counts, scRNA-seq data analysis typically goes through the following steps before clustering: i) normalization (total-count and log-normalization, ii) feature selection, and) iii) dimensional-ity reduction^9, 10^. While normalization adjusts the differences between the samples of individual cells and log normalization reduces the skewness of the data, feature selection seeks to identify the most relevant (top) features (genes) from the large feature space. The top genes are either i) highly variable genes, selected by measuring the coefficient of variation^11, 12^ or ii) highly expressed genes having expression levels higher than the average across all cells^9^.

The performance of downstream analysis, mainly the clustering process is heavily dependent on the quality of selected top features/genes. The typical characteristics of good features/genes are: i) it should encode useful information about the biology of the system ii) should not include features that contain random noise iii) preserve the useful biological structure while reducing the size of the data so as to reduce the computational cost of later steps.

The conventional approach of gene selection based on high variability in the very first stage of the downstream analysis is seemingly simple. However there are two major caveats: i) The observed variability of genes depends on the pseudo-count, which is arbitrary and can introduce biases in the data ii) PCA dimensionality reduction implicitly depends on Euclidean geometry which may not be appropriate for highly sparse and skewed scRNA-seq data. To take care of (i) adding a small pseudocount to all normalized counts prior to log normalization is a common practice in this pipeline. It is required because CPM (counts per million) cannot change the dominating zeros in the scRNA-Seq data, and without that log transformation is not possible. For (2) application of PCA, despite its fast and memory-efficient behavior, is questionable because of the high sparsity, and discrete and skewed nature of the datasets.

In this paper, we address these two challenges, both in themselves, and in their combination. We first present a method that finds the most informative features/genes from the scRNA-seq datasets based on a generalized and wide spectrum entropy measures (Rényi and Tsallis entropy). Although information entropy-based feature selection is a year-long and highly developed subject in the domain of feature selection, applications of entropy in the single-cell domain remains unexplored. There are off course some works exist (e.g^13, 14^), but these are not focused on the informative gene selection in single cell data, instead, Liu et al.,^13^ aim to purify the cluster annotation step whereas Tischendorf et al.,^14^ focuses to quantify differentiation potential in the cells. Here, we indeed demonstrate that employing Renyi and Tsallis entropy in the gene filtering process introduces major advantages both in terms of clustering accuracy, and in terms of a biologically meaningful interpretation (a.k.a. marker selection) of the results. The latter point is established because we are able to reveal marker genes for different cell types for which markers had not been determined through earlier studies. Note that biological marker selection is usually a crucial step in the downstream analysis and is depends on the purity of the cell clusters annotated in the previous stage. Here, we present the most informative gene selection for pure clustering that ultimately leads to a good annotation of the clusters.

Beyond predicting cluster annotation of single cells at utmost accuracy and providing a biologically meaningful interpretation along with it based on scRNA-seq data alone, we also present our method that enables the clustering and annotation in a completely independent data of the same tissue. For this, we split data into a train test ratio of 8:2 and we demonstrate that the selected features in the training set are equally effective in the clustering of the test samples. We also make a comprehensive simulation study to established the proposed method in eight contaminated Gaussian Mixture data. The results prove that our study not only can adopt genes at utmost accuracy, also provides a robust selection with tuned parameters.

**Summary of contributions:** In this work, we provide the following novelties:

1. We provide the first entropy-based gene selection approach for clustering single-cell data. We utilized Renyi and Tsallis entropy that has major advantages over the Shannon method for their controlling parameter (*q*), which makes them less sensitive (robust) against different noises present in the data.
2. Our approach is the first one to explicitly address how to learn the feature relevancy and redundancy using Renyi and Tsallis entropy in the single-cell expression data. We raised an objective function that will minimize conditional entropy between the selected features and maximize the conditional entropy between the class label and feature.
3. Our framework (with tuned parameter) can able to cluster unseen scRNA-seq expression data with utmost accuracy. Clustering hitherto unclassified data is crucial for the annotation of cell types. We present our framework to be effective in this case. We demonstrate that the selected features from the training set are equally valid for the completely unseen test data of the same tissue.
4. Here we derived a new risk factor for Renyi (*R*_*q*_) and Tsallis 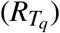 entropy (see Method). We theoretically proved that the iterative selection of features eventually minimizes the Renyi (*R*_*q*_) and Tsallis 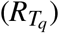 risk, which strengthens the robustness of our objective function.
5. Our method is less sensitive (robust) against different noises present in the data. This is because of the Renyi and Tsallis entropy which has the advantage of controlling their parameters over the Shannon method.
6. It is difficult to achieve good clustering results in small sample single-cell RNA-seq data. Our method also provides good results for small-sample and large-feature sized single cell data. Thus our method can be utilized as a generalized framework for downstream analysis of the single-cell analysis pipeline.

### A short description of feature selection and related works

Selection of relevant features remains an year-long important problem in machine learning domain, because of the ever-increasing size of the dataset. Examples of such high-dimensional data include genomic data, text data, images data, etc, where feature selection plays an important role^15^. The principle motivation of feature selection is to reduce the dimension of large datasets to decrease computational cost and enhance the accuracy of algorithms^16^ applied on the data. Several machine learning and data mining algorithms^17^ are widely used to extract or select meaningful and informative features in different domains such as text categorization^18^, image retrieval, intrusion detection^19^, genomic analysis^20^ and so on. Feature selection algorithms are broadly classified into three categories: the *filter* model^17^, the *wrapper* model^21^ and the *embedded* model^22^.

Filter models are usually based on data filtering and do not include any learning algorithm. Features are selected based on their scores in different statistical tests for their correlation with the outcome variable. So, the features are considered independently and thus ignores the dependencies among them. Despite this disadvantage filter methods are popular in the preprocessing steps because of their simplicity and easy implementation.

A wrapper model requires a learning algorithm to judge the goodness of a subset of features. It searches the non-redundant features to improve the performance of the learning algorithm and hence requires more computational cost than filter methods. Finally, the embedded model takes advantage of both the wrapper and the filter methods to select a subset of features.

Many interesting feature selection approaches were proposed in earlier studies, they are-recursive feature elimination^23^, sequential feature selection algorithms^24^, genetic algorithms based feature selection approaches^25^, mutual information or entropy based feature selection^26^, etc.

Several entropy based filter methods have been proposed in the literature^27^. Most of these methods use Shannon’s entropy. Renyi and Tsallis entropy^28^ is another option that needs to be investigated in this context. Recently, some advances in the fields of security and privacy have reinvigorated interest in Renyi and Tsallis entropy^29^ and thus they are ripe for application in biological data as well.

There exist different works on feature selection based on entropy measures. In^30^, a novel algorithm called MIFS was proposed to automatically estimate the effective number of features using mutual information (Shannon Entropy). Jiang et al.,^31^ proposed a new model of relative decision entropy for feature selection, which is an extension of Shannon’s information entropy in rough sets. A study was conducted by Lopes et al.^32^ defining the sensitivity of entropy methods which is applied to a well-defined problem of gene regulatory network.^33^ proposed a new information-theoretical method for feature selection using Renyi min-entropy. Another entropy based feature selection method is proposed in^27^ for text categorization.

## Results

In the following, we will first describe the workflow of our analysis pipeline and the basic ideas that support it.

First, we utilized a basic framework for preprocessing the scRNA-seq data. We use recent filtering techniques to filter out genes and cells from the data and use this as an input of the proposed method.

We then carry out a simulation study on contamination Gaussian Mixture data that proves that our study can adopt genes at the utmost accuracy. The study also provides a robust selection using the tuned parameter, *q*.

Subsequently, in our real experiments, we choose genes in five single cell RNA sequence data. Corroborated by our simulations, a sufficient number of informative genes are selected, which is potentially actionable in single cell data analysis in less computational cost.

Finally, our study yields the selected genes in the frame of marker genes in literature, documenting the plausibility of our predictions.

### Workflow

The figure 1 describes the workflow of our analysis pipeline. All important steps are discussed in the paragraphs of this subsection.

**Figure 1.**
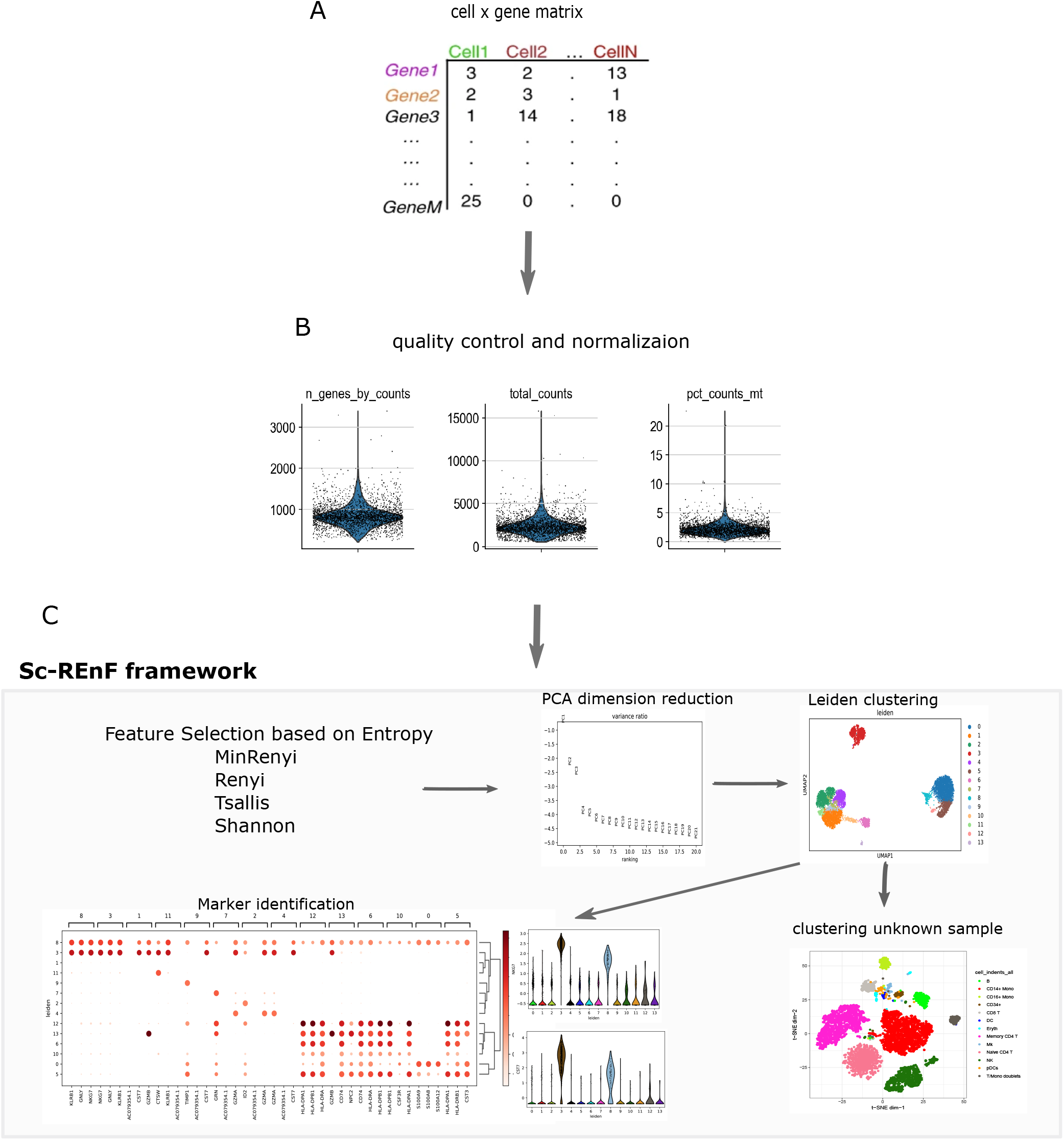
A brief framework of our study: Panel-A and B: scRAN-seq count matrix are downloaded and preprocessed (quality control and normalization). sc-REnF is applied for gene/feature selection using MinRenyi, Renyi, Tsallis and Shannon entropy measure (panel-C (1)). Dimension reduction and clustering is performed to group the cells in clusters (Panel-C (3)). Marker gene analysis and clustering of unknown samples is performed using the results of panel-C (1)

#### Preprocessing of Raw Datasets

Single-cell RNA sequence raw datasets are downloaded from publicly available sources. The RNA counts are organised as a matrix *M*_*cl*×*ge*_, where *cl* is the number of cells and *ge* is the number of genes. Each element [*M*]_*ij*_ represents count of the *i*^*th*^ cell in the *j*^*th*^ gene. If more than a thousand of genes are expressed (non zero values) in one cell, then the cell is termed as good. We assume one gene is expressed if the minimum read count of it exceeds 5 in at least 10% of the good cells. The data matrix *M* with expressed genes and good cells is normalized using a linear model and normality based normalizing transformation method (Linnorm)^34^. The resulting matrix is then *log*_2_ transformed by adding one as a pseudo count. Thus, the preprocessed matrix *M*_cl′×*ge′*_ is derived and is used in the entropy based feature selection model.

#### Entropy based model for Feature Selection

The preprocessed data matrix *M′* is used in the proposed entropy based (Renyi and Tsallis) feature selection models. First, a feature ranking is performed based on the relevancy between all features and class labels (see equation 14 in Method). The top rank (most relevant) feature is selected based on the relevancy score. Then redundancy of remaining features with the selected one is computed next (see equation 15 in Method). The process includes a minimum redundant feature in the selected features list. The process goes in an iterative way by adding the most non-redundant features in each step.

#### Dimensionality reduction and clustering

Here we use principal component analysis to reduce the dimension of the data before clustering. We adopt the conventional process used in the Scanpy^35^ package for dimensionality reduction and clustering process. We pick the top 15 PCs and create a neighborhood graph of cells using the PCA representation. It is assumed that the top PCs are likely to represent the biological signals while the latter PCs are adopting the noise in the data. The dominant factors of heterogeneity are likely to be captured by the top PCs. The neighborhood graph of cells is then directly clustered by Leiden graph-clustering method^36^ to groups cells into different clusters.

#### Marker identification

We compute ranking for the highly differential genes within each cluster. Here we identify DE genes in each cluster using the Wilcoxon Ranksum test, but other statistical tests may be utilized otherwise. We select the top ten DE genes from each cluster based on the p-values.

#### Clustering of unknown samples

For cells of the unknown type, our method can able to cluster with the selected genes, yielding a good clustering that is as fine-grained as is justified by the available true labeled data. Clustering unknown samples are crucial in scRNA-seq data analysis and can be addressed by a supervised or unsupervised way. In supervised technique, the model trained with reference data can be used to predict the cell types of samples. In our case (which is supervised), we observed that the selected genes in reference data can be useful to cluster the unknown samples of the same tissue. The clusters may be annotated by matching the DE genes with canonical markers. This provides the key element of our approach to work in practice.

#### Validation of selected features/genes

We validate our selected features/genes in several ways. First, a comprehensive simulation study is conducted to ensure the robustness of our proposed method. Clustering results on synthetic data are evaluated using the Adjusted Rand Index (ARI) score. To validate the proposed risk measure, a stability test has also been conducted. Second, the clustering results on scRNa-seq data are evaluated using ARI to ensure proper and accurate partitioning of cells. Having a good partitioning, thirdly we compute DE genes within each cluster which are actually treated as marker genes of specific cells. Finally, clustering on unknown samples is validated through a test set built from labeled scRNA-seq data. TSNE 2 visualization is used in all cases to visualize the data with original and predicted annotation.

### Clustering Synthetic Data: Validation

Clustering performance on synthetic contaminated Gaussian mixture datasets (see Method) is evaluated using Adjusted Rand Index (ARI). The ARI score ensures the quantity of matching between predicted cluster labels with known groups and is ranging from 0 (when clustering prediction is random) to 1 (when clustering is perfectly coherent with the known labels)^37^. *k*–means clustering is utilized to cluster the synthetic Gaussian mixture dataset. Renyi and Tsallis entropy with twelve q values ranging from 0.1 − 7 are used to select features from the synthetic datasets (both for overlapping and non-overlapping case, see method for details). We make a note of q values for which the two methods perform well. It can be observed from the Figure 2 that for the Renyi entropy ARI achieves high score in the range [0.3,1.5] of q-values. Similarly, for Tsallis entropy the range of q-values for which ARI scores is high are [0.3, 1.3]. All results are computed for the synthetic data containing four overlapped and four non-overlapped clusters. The selected range of *q*-values are utilized in real life scRNA-seq data for feature selection, and *q*–value yields highest ARI, is reported in the table 1.

**Figure 2.**
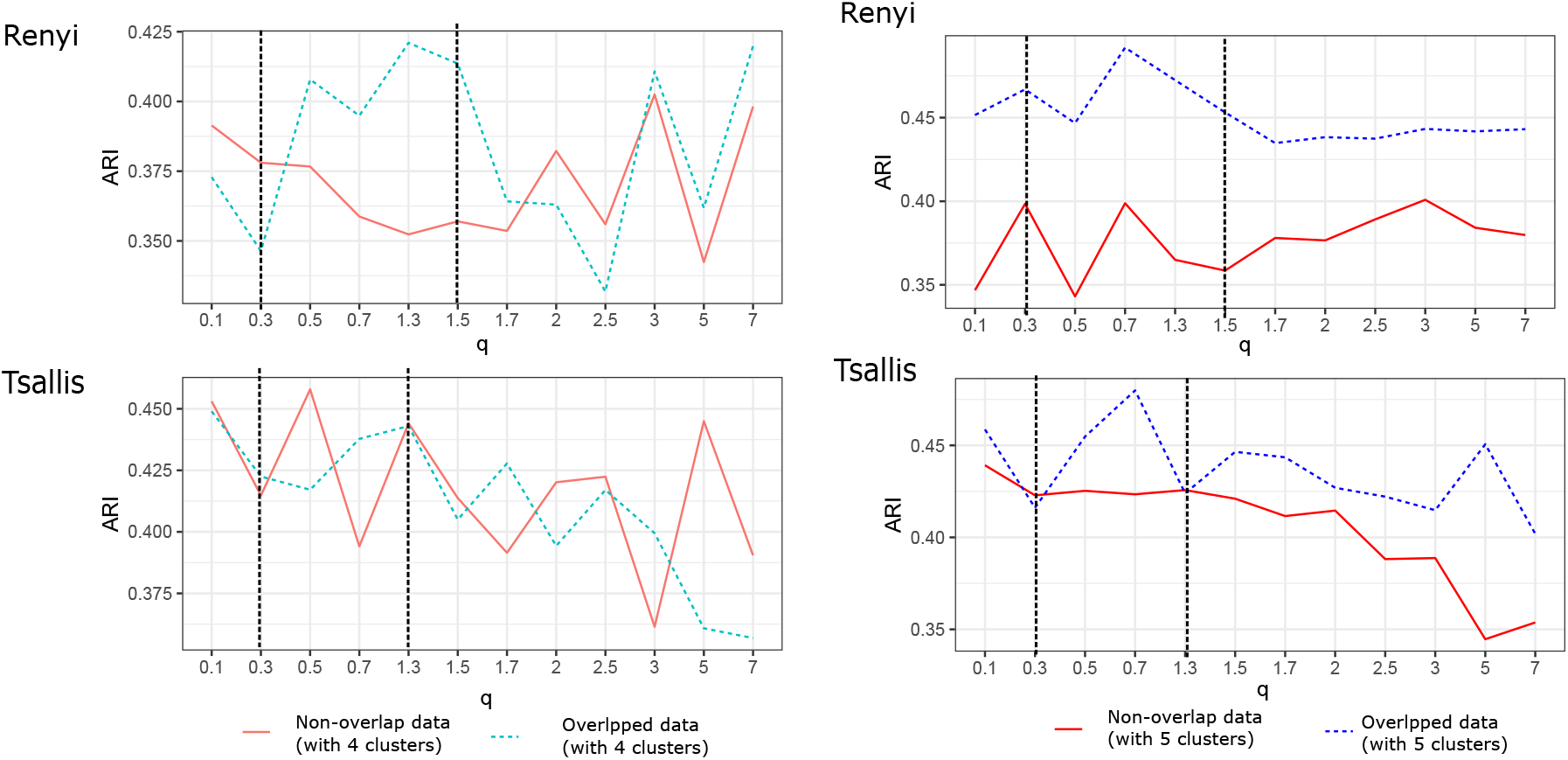
ARI scores for clustering of synthetic data containing four overlapped and non-overlapped clusters. sc-REnF is used with 12 *q–values* for Renyi and Tsallis entropy within the range [0.1,7].

**Table 1.** Adjusted Rank Index measured on the clustering results on scRNA-seq datasets

| Serial # | Datset | Min Renyi | Renyi (0.7) | Tsallis (0.3) | Shannon | Cluster Number (#k) |
| --- | --- | --- | --- | --- | --- | --- |
| 1 | CBMC | 0.56 | 0.63 | 0.70 | 0.55 | 14 |
| 2 | Darmanis | 0.37 | 0.41 | 0.45 | 0.40 | 8 |
| 3 | Yan | 0.72 | 0.87 | 0.66 | 0.69 | 4 |
| 4 | Pollen | 0.89 | 0.93 | 0.61 | 0.56 | 11 |

### Clustering scRNA-seq Data: Validation

After knowing the range of *q* for the Renyi and Tsallis method we applied sc-REnF on four scRNA-seq datasets. Clustering is done for the sake of validation. The selected genes by sc-REnF are used for the downstream analysis (PCA and clustering) and ARI score is computed after clustering. Table 1 illustrates the clustering performance on the four scRNA-seq datasets using Min Renyi, Renyi, Tsallis, and Shannon methods. It is observed that sc-REnF responds well for Renyi and Tsallis measures compare to the other two methods. Renyi based method achieves the highest ARI score for Pollen dataset *q*- value of 0.7. Similarly, the Tsallis method yields the highest ARI score for the darmanis dataset with *q*-value of 0.3.

After gene selection, we perform dimensionality reduction using traditional PCA and create a neighborhood graph using the PC components, which is then clustered by the Leiden clustering method.

#### Clustering results on Darmanis data

Figure 3 shows the results of applying sc-REnF on Darmanis data. To explore how well sc-REnF can selects genes from Darmanis data we use a clustering technique to group cells into the cluster and match them with the original level. After gene selection, we use PCA for dimensionality reduction and Leiden clustering for grouping of cells. Figure 3 panel-A shows the t-SNE plot of 466 cells with the original level. The Leiden clustering produces eight clusters (shown in panel-B) which are then match with the original level. Figure 3 Panel-C represents the percentage of matched samples between the resulting clusters and the original level. As can be seen from Figure 3 panel-C in most of the cases, one cluster can be determined by a unique cell type (e.g. cluster-1 is matched with fetal quiescent cluster-6 is matched with oligodendrocytes and so on). Panel-D shows the cells of a particular type with the identified matched clusters (color-coded). Although Panel-C can visualize the matching behavior between identified clusters and original cell labels, Panel-D explores the matching of cell type at the individual level. It can be noticed that for cell type ‘OPC’ and ‘microglia’ are in the same cluster (cluster-5), whereas for ‘astrocytes’ most of the cells are going to cluster-4. There are off course some cells (e.g. ‘neurons’, ‘hybrids’) exist which go to several clusters, nevertheless, one particular cluster contains the majority of the cells, suggesting a good clustering of the data.

**Figure 3.**
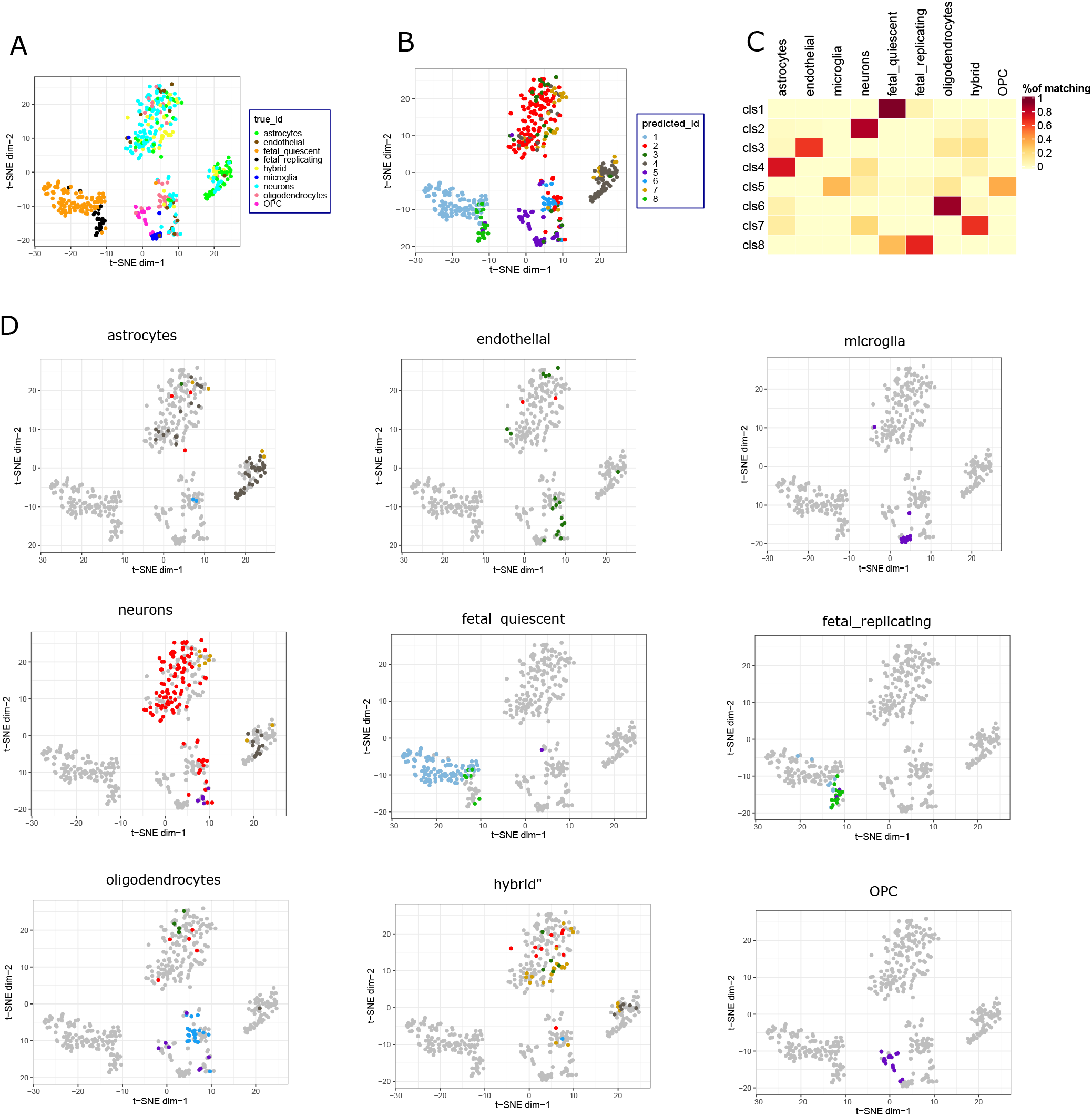
Clustering results of Darmanis data after gene selection. Panel-A and -B represents t-SNE visualization of data with original and predicted cluster label respectively. Panel-C shows a heatmap that represents percentage of matching samples between eight identified cluster and different cell types. Panel-D depicts visualization of samples coming from individual cells and their corresponding predicted clusters (color coded)

#### Clustering results on CBMC data

Application of sc-REnF on 8000 cord blood mononuclear cells (CBMCs) produces 14 clusters. Figure 5 panel-A and B shows the t-SNE visualization of cells with original labels and predicted cluster labels, respectively. Most of the clusters such as cluster-2, cluster-4, cluster-14 determine unique cells in the data. For example, cluster-2 captures most of the samples of CD14+ Mono cells, while cluster-4 and cluster-14 represent samples of ‘Nk’ (Natural killer) and erythrocyte cells. Some clusters represent more than one cell, such as cluster-13 includes DCs and pDCs, cluster-11 includes erythrocyte, Mks, and CD34+ cells. Individual mapping of cells to different clusters is depicted in Figure 5, panel-D. For example ‘NK’ cells are captured by two clusters, cluster-4 and cluster-12. Despite being multiple associations of clusters into one particular cell type, most of the cases one cluster captures major samples of a particular cell type.

### Clustering unknown sample: Validation through test data

Clustering unknown sample is crucial for scRNA-seq analysis pipeline. Here we addressed this by performing a cross validation with a train test split of data in the ratio 7:3. First, the training dataset is used in sc-REnF to select informative genes. Top 50 genes are selected for clustering and validation. The performance of sc-REnF is computed on the test data using the top selected genes in the earlier step. We performed clustering on test data with the selected genes and ARI value is reported. The experiment is repeated 20 times with a random split of train-test data (7:3 ratio) in each case. Table 2 shows the median and standard deviation of the ARI score for Renyi and Tsallis measure.

**Table 2.**
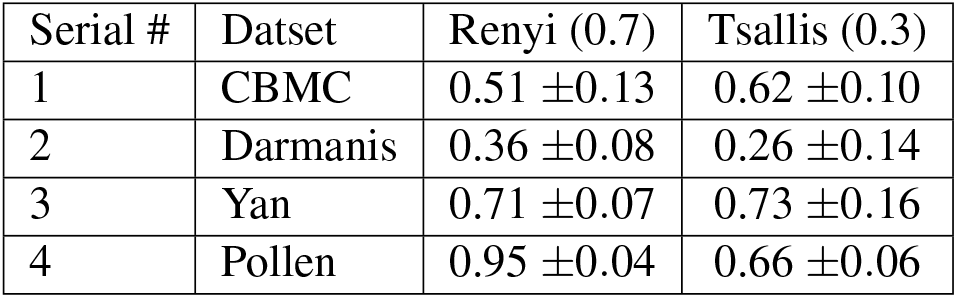
Adjusted Rank Index measured on the clustering results on unknown test samples

| Serial # | Datset | Renyi (0.7) | Tsallis (0.3) |
| --- | --- | --- | --- |
| 1 | CBMC | 0.51 ±0.13 | 0.62 ±0.10 |
| 2 | Darmanis | 0.36 ±0.08 | 0.26 ±0.14 |
| 3 | Yan | 0.71 ±0.07 | 0.73 ±0.16 |
| 4 | Pollen | 0.95 ±0.04 | 0.66 ±0.06 |

### Marker Gene Selection

The clustering results are further utilized in the marker gene selection for different cell types. From each cluster DE genes are identified that drives the separation between clusters. Here we have used Wilcoxon rank-sum test to directly assesses separation between the expression distributions of different clusters. For darmanis dataset, Figure 4 panel-A–D and Figure 6 panel-A–E represent the results and visualizations of marker gene analysis in darmanis and CBMC data, respectively. The higher expression values of top five DE genes for a particular cluster (see Figure 4 panel-B and Figure 6 panel-C) represents the presence of marker genes within the selected gene sets. The result is clearly visible in the dotplot of average expression values of top DE genes shown in Figure 4 panel–C and Figure 6 panel-E. We manually match the identified DE genes with a cell marker database published by Zhang et al.^38^ and report the matched marker in table 3 for darmanis data. For Darmanis data the DE genes which are not included in the cell marker database and keep a higher expression in some particular cluster may be treated as a new marker of cell represented by that cluster. For example gene ‘VIP’ and ‘PTPRS’ shows higher expression in cluster-3 and cluster-7, respectively (shown in Figure 4, panel-D), suggesting a suitable candidate for marker gene of cell astrocytes and fetal replicating cells (see Figure 4, panel-C for the cell annotation of cluster-3 and cluster-7). For CBMC data, gene ‘S100A9’, ‘NKG7’ ‘LST1’ have higher expression in cluster-0, cluster-3 and cluster-5, respectively, suggesting novel markers for cell Naive CD4T, NK and CD14+Mono cells (see Figure 6, panel-C for the cell annotation of cluster-0, cluster-3 and cluster-5).

**Figure 4.**
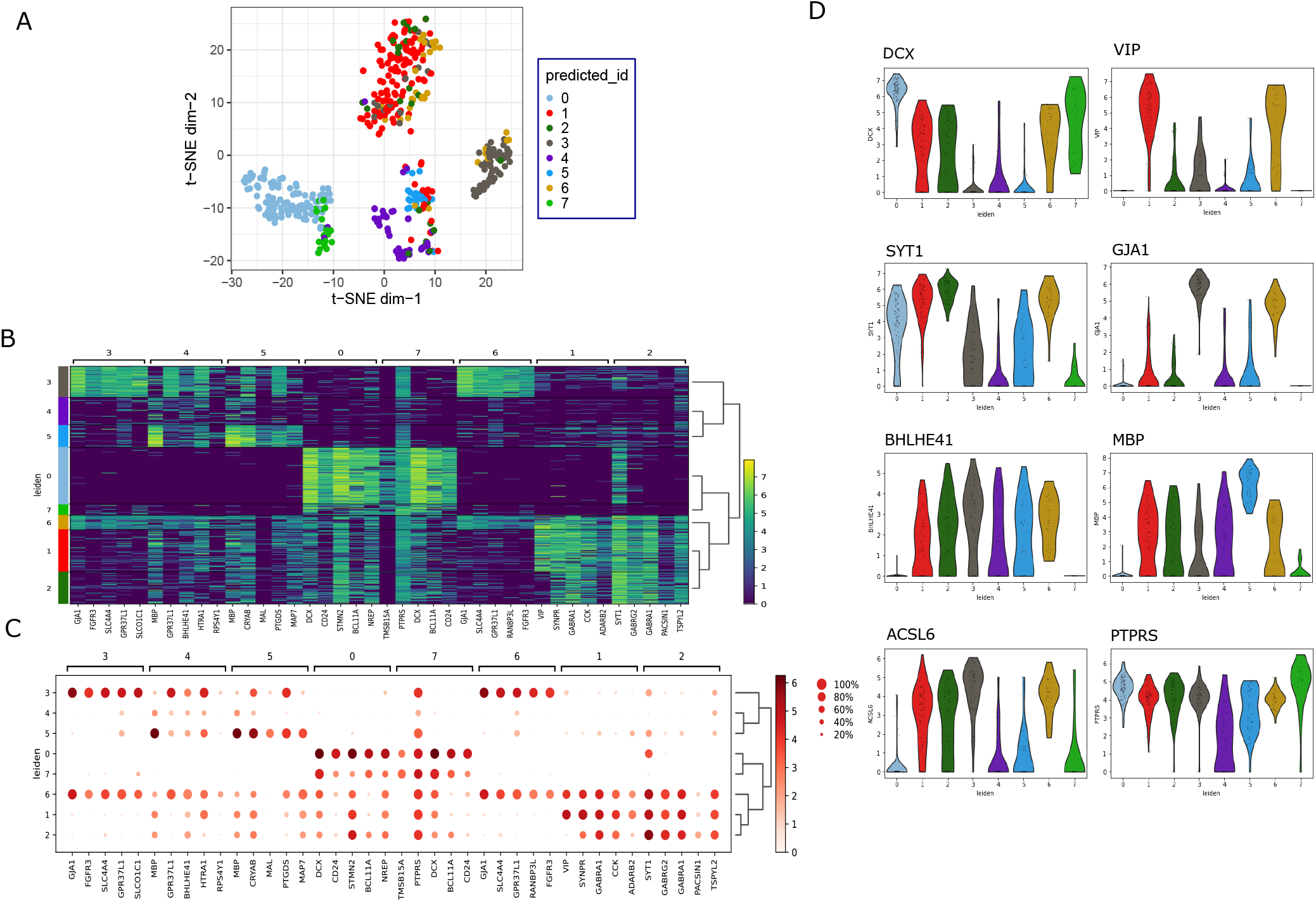
Results of marker gene analysis in Darmanis data. Panel-A shows t-SNE visualization of data with predicted cluster labels. Panel-B shows heatmap of expression values of top five DE genes in each cluster. Panel-C shows the average expression of top five DE genes within each cluster. Panel-D represents expression of identified marker genes across different clusters.

**Figure 5.**
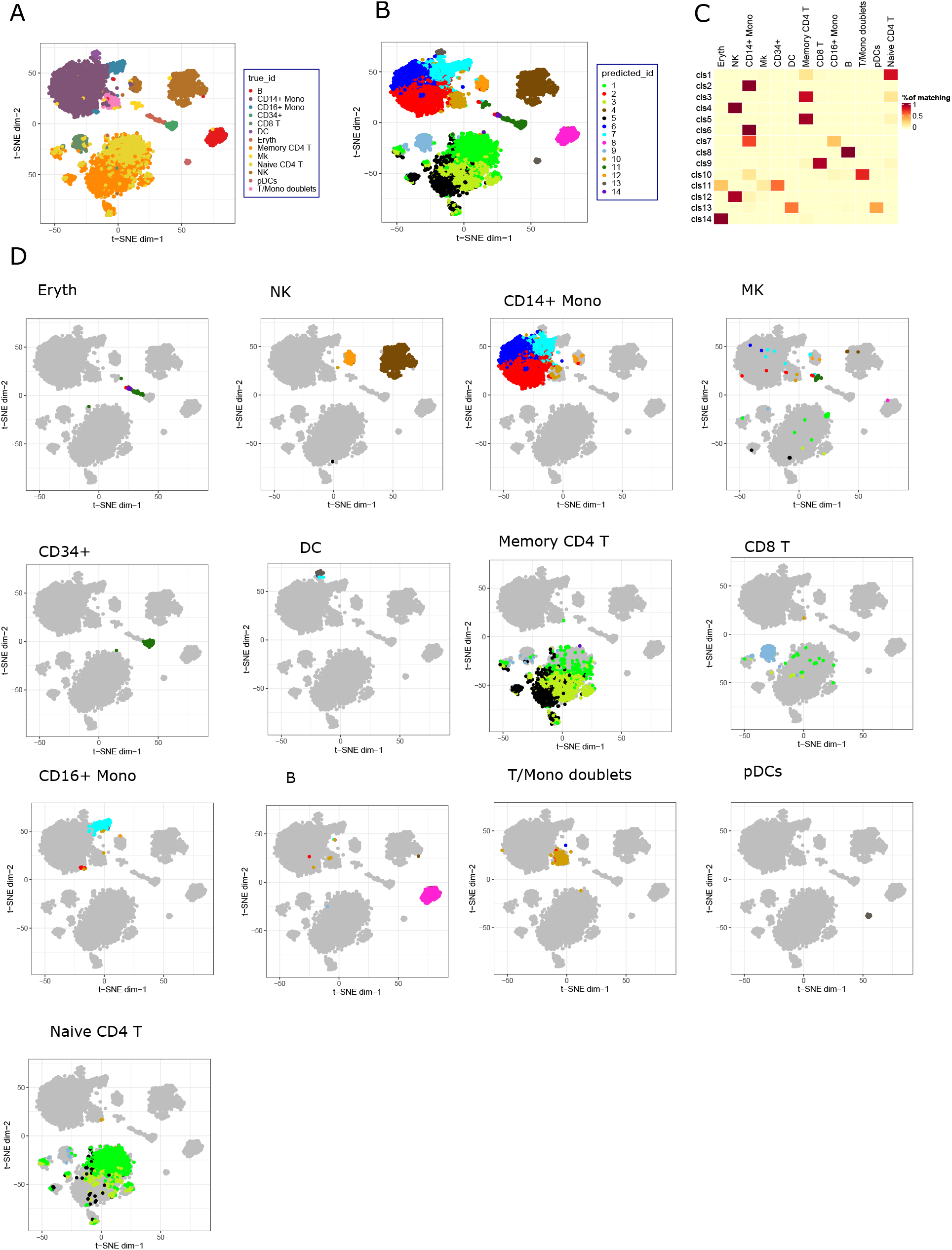
Clustering results of CBMC data after gene selection. Panel-A and -B represents t-SNE visualization of data with original and predicted cluster label respectively. Panel-C shows a heatmap that represents the percentage of matching samples between 14 identified cluster and 13 different cell types. Panel-D depicts visualization of samples coming from different immune cells and their corresponding predicted clusters (color coded)

**Figure 6.**
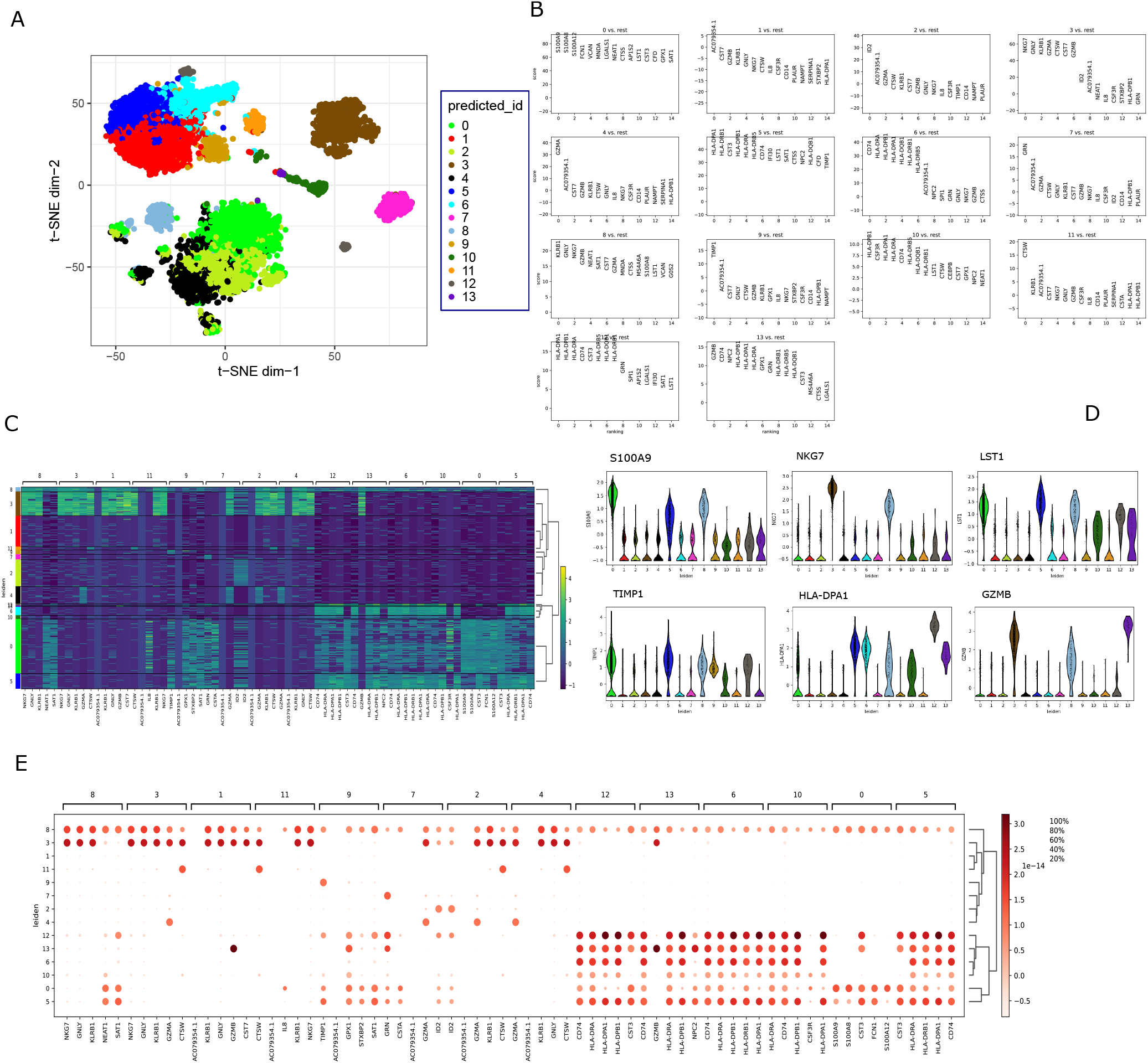
Results of marker gene analysis in CBMC data. Panel-A shows t-SNE visualization of data with predicted cluster labels. Panel-B shows ranking of genes in different cluster using Wilcoxon rank-sum test. Panel-C shows heatmap of expression values of top five DE genes in each cluster. Panel-D shows the average expression of top five DE genes within each cluster. Panel-E represents expression of identified marker genes across different clusters

**Table 3.** Table shows the top 10 marker genes for Darmanis data. Some marker genes are found in the cellmarker database^38^ (shown in bold text). The genes are shown with their respective p-values (Wilcoxon Rank-sum test).

| # Serial | Cluster Number & P-value |  |  |  |  |  |  |  |
| --- | --- | --- | --- | --- | --- | --- | --- | --- |
|  | Cls0 | Wilcox_Pval | Cls1 | Wilcox_Pval | Cls2 | Wilcox_Pval | Cls3 | Wilcox_Pval |
| 1 | **DCX** | 7.23E-51 | VIP | 4.52E-36 | **SYT1** | 1.66E-22 | **GJA1** | 7.69E-35 |
| 2 | **CD24** | 6.09E-48 | SYNPR | 9.24E-34 | GABRG2 | 4.60E-21 | **FGFR3** | 1.09E-33 |
| 3 | **STMN2** | 1.34E-46 | GABRA1 | 3.92E-26 | GABRA1 | 1.02E-14 | **SLC4A4** | 3.99E-32 |
| 4 | **BCL11A** | 1.97E-42 | CCK | 1.66E-25 | PACSIN1 | 3.70E-13 | **GPR37L1** | 1.02E-31 |
| 5 | NREP | 4.01E-41 | ADARB2 | 5.88E-21 | TSPYL2 | 5.84E-11 | **SLCO1C1** | 1.20E-31 |
| 6 | **TMSB15A** | 6.20E-31 | OSBPL1A | 1.79E-19 | SPOCK2 | 2.16E-10 | RANBP3L | 2.12E-29 |
| 7 | FNBP1L | 9.29E-24 | GABRG2 | 1.59E-18 | CCK | 1.15E-09 | **PRODH** | 1.31E-27 |
| 8 | PTPRS | 3.04E-15 | TSPYL2 | 3.53E-15 | ENTPD3 | 1.46E-09 | COL5A3 | 1.40E-26 |
| 9 | **RBFOX2** | 6.44E-10 | MYT1L | 6.88E-14 | **LGI1** | 2.85E-07 | GABRA2 | 3.25E-26 |
| 10 | MYT1L | 7.12E-08 | SPOCK2 | 3.00E-13 | MYT1L | 9.00E-07 | **MLC1** | 3.55E-26 |
| # Serial | Cluster Number & P-value |  |  |  |  |  |  |  |
|  | Cls4 | Wilcox_Pval | Cls5 | Wilcox_Pval | Cls6 | Wilcox_Pval | Cls7 | Wilcox_Pval |
| 1 | **MBP** | 0.098026 | **MBP** | 7.10E-25 | **GJA1** | 7.47E-11 | **TMSB15A** | 5.58E-08 |
| 2 | **GPR37L1** | 0.390515 | **CRYAB** | 3.51E-23 | **SLC4A4** | 8.35E-10 | PTPRS | 6.20E-06 |
| 3 | **BHLHE41** | 0.418991 | **MAL** | 3.08E-19 | **GPR37L1** | 2.32E-09 | **DCX** | 0.003346 |
| 4 | **HTRA1** | 0.615749 | PTGDS | 1.88E-15 | RANBP3L | 9.11E-09 | **BCL11A** | 0.02091 |
| 5 | RPS4Y1 | 0.346481 | MAP7 | 3.35E-15 | **FGFR3** | 9.92E-09 | **CD24** | 0.079551 |
| 6 | MFAP3L | 0.333967 | CAMK2N1 | 3.11E-06 | VIP | 2.79E-08 | **MLC1** | 0.104821 |
| 7 | TSPYL2 | 0.286378 | **HTRA1** | 8.91E-05 | **SLCO1C1** | 4.57E-08 | FNBP1L | 0.141708 |
| 8 | RSPH9 | 0.183538 | OSBPL1A | 0.000326 | **ACSL6** | 1.27E-07 | **SLCO1C1** | 0.188929 |
| 9 | **MAL** | 0.108362 | MFAP3L | 0.001364 | SPOCK2 | 1.34E-07 | NREP | 0.570809 |
| 10 | **SLC14A1** | 0.046909 | ADARB2 | 0.010814 | SYNPR | 2.58E-07 | RSPH9 | 0.846317 |

### Stability checking of sc-REnF

Here we explore an additional advantage of sc-REnF over the other measures by validating its stability of performance. A non-parametric statistical test KruskalWallis Test^39^ is utilized to examine the stability of ARI scores resulted from the clustering results. We vary the number of features from range 10 to 50 and for each case, we compute the ARI score after clustering. Thus for one method (e.g. Renyi) and for one dataset, we get five ARI scores (for #feature=10, 20,30,40,50) representing the clustering performance with different selected features. To know the variation of ARI scores across all the datasets for a particular method (e.g. Renyi), we performed KruskalWallis Test. ARI scores are computed 50 times for each method, and the median of the scores is given to the KruskalWallis test. Table 4 shows the result (p-values and chi-squared values) of the KruskalWallis test. Although all the methods produce a stable results with low p-values, nevertheless the Tsallis (with q-value=0.7) and Renyi (with q-value=0.3) show more stable performance among other methods. These results may be treated as a straightforward implication of the theoretical proof present in n.

**Table 4.**
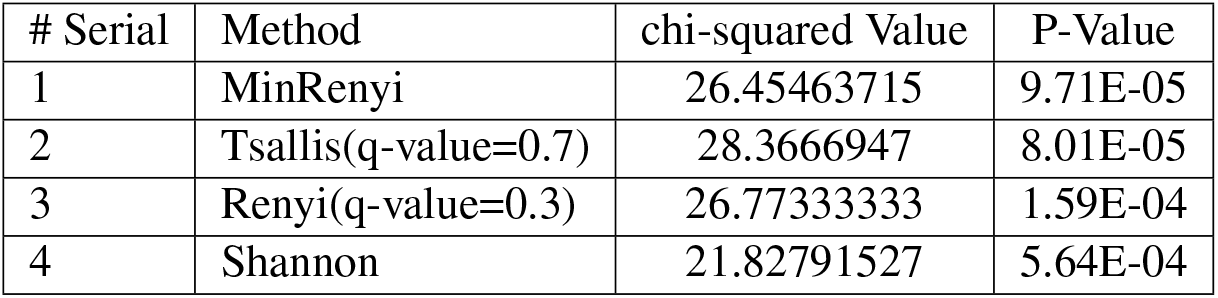
Stability performance of sc-REnF: p-values is reported on the basis of Kruskal-Wallis test on the ARI scores obtained from clustering results of synthetic data

## Discussion

Clustering of cells in scRNA-seq data is an essential step for cell type discovery from a large population of cells. Owing to the large feature/gene set of scRNA-seq data, selection of most variable genes are crucial in the preprocessing step, which has immense effect in the later stage of downstream analysis. The proposed method sc-REnF addressed this issue by using an entropy (Renyi, Tsallis) based feature selection method for identifying possible informative genes in the preprocessing steps. sc-REnF has the advantage over the conventional statistical approach that it can consider the cell-to-cell dependency based on generalized and wide spectrum entropy measures Renyi and Tsallis. We demonstrated that sc-REnF using Renyi and Tsallis method introduces major advantages both in terms of clustering accuracy and in terms of marker gene detection in the downstream analysis of scRNA-seq data.

sc-REnF yields a stable feature/gene selection with a controlling parameter (*q*) for Renyi and Tsallis entropy. The optimal controlling parameter (*q*) is determined by applying it in a synthetic contaminated Gaussian mixture dataset. We later demonstrated that the range of selected *q* – *values* is applicable in the real life scRNA-seq data clustering task. The four scRNA-seq data where we apply sc-REnF yields accurate clustering results which are validated by the ARI index. The stability of sc-REnF is demonstrated by evaluating the performance of it using KruskalWallis test. While applying sc-REnF multiple times with varying number features, the resulting ARI scores employ a minimum deviation (p-value « 0.05 for Kruskal-Wallis) for Renyi and Tsallis entropy.

Although the primary objective of sc-REnF is variable gene selection in the preprocessing step of scRNA-seq data analysis, we extend the process towards the later stage of downstream analysis. We employ clustering technique to groups the cells using those selected genes. A precise clustering of cells also demonstrates the efficacy of our method for selecting the most variable genes in the first stage. This facilitates the selection of novel marker genes within each cluster. We pinpoint several markers, which shows a high expression level within a particular cluster, among them some of are also identified in previously published cell marker database.

Clustering of unknown samples based on the reference data is a crucial problem for the identification of cell types in scRNA-seq cell classification. We addressed the problem by cluster the unknown samples using the selected genes in the reference data. We demonstrate the advantage of using selected genes by sc-REnF in clustering of the unknown test sample. We observed good ARI scores in the clustering of test samples, suggesting the selected genes from the reference data is also effective to produce a perfect clustering in a completely unknown test sample.

The execution time of sc-REnF is directly proportional to the number of selected features and can be expensive when one needs to select a large number of features. However, this can be easily tackled with ever-increasing computing power in advanced servers. Additionally, the regularization parameter has not been considered in our proposed approach, which may sometime make the algorithm susceptible to overfitting unless carefully employed.

Taken together, the proposed method sc-REnF not only has good performance on informative gene selection in the preprocessing step but also has the ability to explore the classification of unknown cells in the scRNA-seq data. Despite being applied in feature selection of different domains, the application of Renyi and Tsallis entropy shows good potential in gene selection and cell clustering of scRNA-seq data. Results show that sc-REnF not only leads in the domain of robust feature (gene) selection analyses but accelerate the investigations of cell type definition in large scRNA-seq data as well. We believe that sc-REnF may be an important tool for computational biologists to explore the most informative genes and marker genes in the downstream analysis of scRNA-seq data.

## Methods

### Overview of Datasets

#### Synthetic Gaussian Mixture Data

We generated eight synthetic Gaussian mixture datasets (overlapping and non-overlapping) having *k* = {2, 3, 4, 5} number of clusters. Each data contains 50 relevant and 250 irrelevant features with 500 samples.

The relevant features are generated by varying the mean *μ* of the dataset in the range [−5, 20] for non-overlapping and in the range [−2, 3] for overlapping with a fixed covariance matrix Σ in each case. The covariance matrices (Σ) are computed as: Σ = (*ρ*^|*i*−*j*|^), where *i, j* are the row and column index of the covariance matrix, *ρ* = 0.5.

For generating the irrelevant feature white Gaussian noise^13^ is generated and added to the constructed synthetic datasets. The R package *Add.Gaussian.noise* is utilized to generate Gaussian noise with mean 0, and standard deviation 1. A detailed descriptions of the synthetic data is given in the table 5, and in table 6. Figure 7 depicts a 2-dimensional tSNE visualization of the generated synthetic gaussian mixture datasets. To make the data noisy we add contaminated noise in the constructed synthetic data by using 8 : 2 mixing ratio among the classes. The following steps are used for contamination of one synthetic data (assuming 2 class *c* and 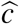 data).

1. 20 percent samples of one class (*c*) (with mean *μ* and covariance matrix Σ) are replaced with samples generated with mean 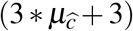 and covariance matrix Σ of other class 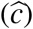 where 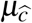 is the mean of the other class 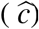.
2. The process is repeated for all eight Gaussian mixture datasets.

**Table 5.**
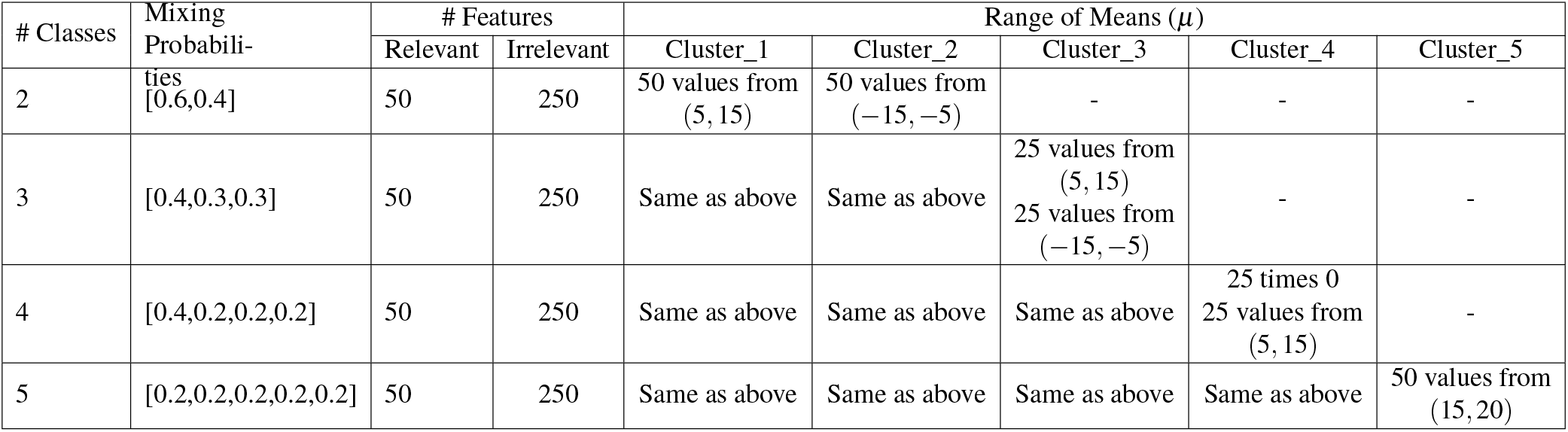
Description of non-overlapping synthetic Gaussian mixture Data

**Table 6.**
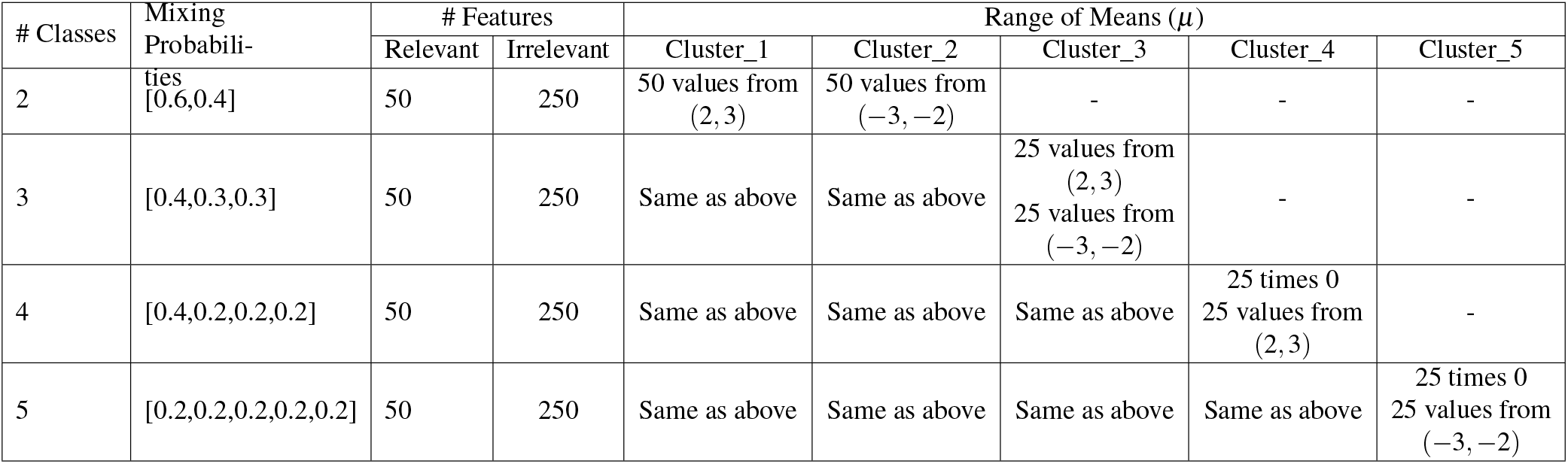
Description of overlapping synthetic Gaussian mixture data

| # Classes | Mixing Probabili- | # Features | | Range of Means ($\mu$) | | | | |
| --- | --- | --- | --- | --- | --- | --- | --- | --- |
|  |  | Relevant | Irrelevant | Cluster_1 | Cluster_2 | Cluster_3 | Cluster_4 | Cluster_5 |
| 2 | ties [0.6,0.4] | 50 | 250 | 50 values from (2, 3) | 50 values from (-3, -2) | - | - | - |
| 3 | [0.4,0.3,0.3] | 50 | 250 | Same as above | Same as above | 25 values from (2, 3)<br>25 values from (-3, -2) | - | - |
| 4 | [0.4,0.2,0.2,0.2] | 50 | 250 | Same as above | Same as above | Same as above | 25 times 0<br>25 values from (2, 3) | - |
| 5 | [0.2,0.2,0.2,0.2,0.2] | 50 | 250 | Same as above | Same as above | Same as above | Same as above | 25 times 0<br>25 values from (-3, -2) |

**Figure 7.**
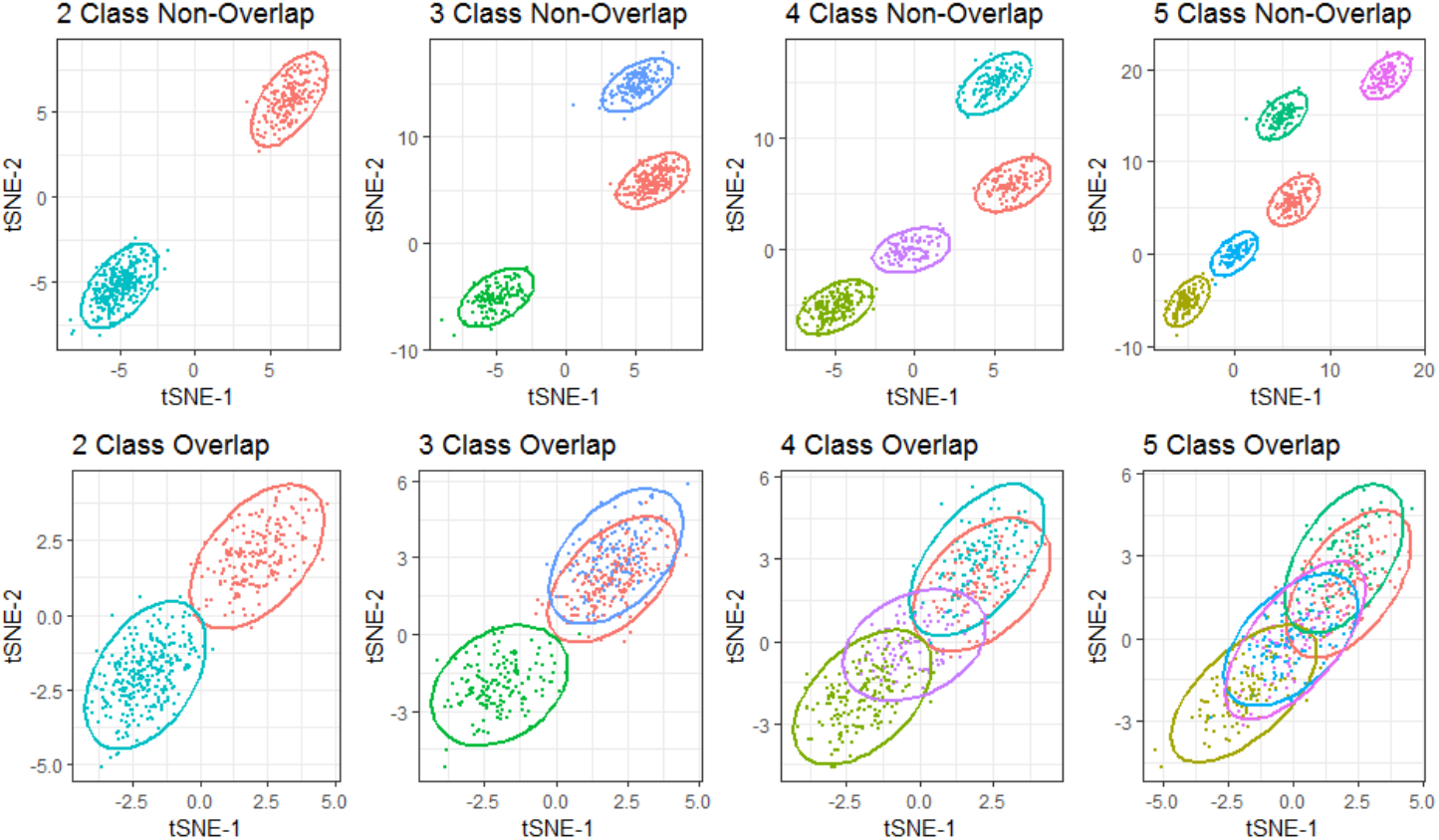
Figure shows tSNE visualization of eight synthetic Gaussian Mixture Datasets. The upper row represents datasets of non-overlapping class whereas the lower row represents the same for overlapping classes.

#### Single Cell RNA sequence dataset

The following single-cell RNA sequence datasets are used for evaluation of the proposed method.

##### Yan

This is a human preimplantation embryo and embryonic stem cell dataset. The average total read count in the expression matrix is 25,228,939 reads. There are 7 cell types, including labelled 4-cell, 8-cell, zygote, Late blastocyst and 16-cell.[GEO under accession no. GSE36552;^40^].

##### Pollen

The data library was generated from 600 individual cells in parallel. It contains 11 cell types. [GEO under accession no GSM1832359;^41^]

##### Darmanis

It contains single cell RNA sequencing on 466 cells to capture the cellular complexity of the adult and fetal human brain at a whole transcriptome level. Healthy adult temporal lobe tissue was obtained from epileptic patients during temporal lobectomy for medically refractory seizures. [GEO under accession no GSE67835;^42^].

##### CBMC

Cord blood mononuclear cells (CBMC) were profiled by CITE-seq which is proposed to measure both cellular protein and mRNA expression in one cell, by using oligonucleotide-labeled antibodies. The data set consists of the expression levels of 2000 mRNAs and 13 protein, individually measured in 8,000 cord blood mononuclear cells (CBMCs).The dataset is available in the GEO website under the accession GSE100866.

The brief description of the dataset is given in Table 7.

**Table 7.**
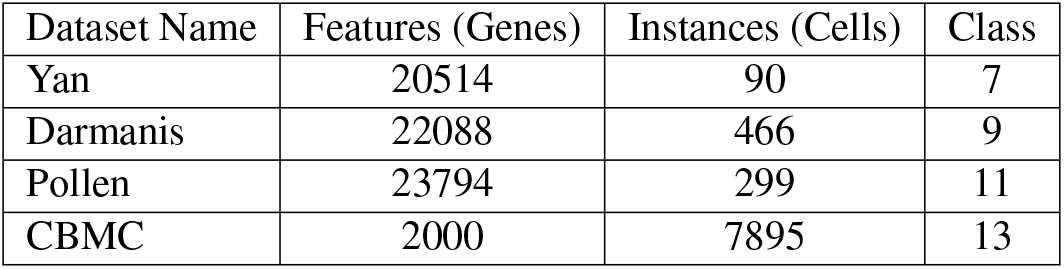
A brief summary of the scRNA datasets used in this study

### Entropy and Risk Functions

For the sake of completeness, let us start with a brief description of different entropy measures, see, e.g.,^43^ for details. Throughout this section, we consider three discrete random variables *X*, *Y* and *Z* having supports {*x*_1_, ..., *x*_*d*_}, {*y*_1_, ..., *y*_*p*_} and {*z*_1_, ..., *z*_*n*_}, respectively. For each *i* = 1, 2, ..., *d*, *j* = 1, 2, ..., *p* and *k* = 1, 2, ..., *n*, let us denote *p_i_* = *P*(*X* = *x*_*i*_), *p*_*ijk*_ = *P*(*X* = *x*_*i*_,*Y* = *y*_*j*_, *Z* = *z*_*k*_), *p*_*i|jk*_ = *P*(*X* = *x*_*i*_|*Y* = *y*_*j*_, *Z* = *z*_*k*_) and so on.

#### Shannon Entropy

For a random variable *X*, the most popular Shannon entropy is defined as

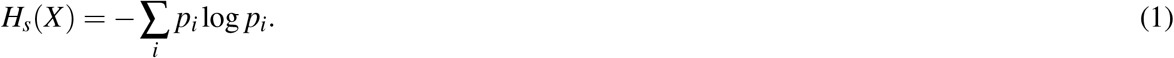

For more than one variables, one can suitably construct the joint or the Renyi entropy measures. For example, the Shannon conditional entropy of the random variable X given two random variables Y and Z is defined as The conditional Shannon entropy of three random variable *X*, *Y*, and *Z* is described below

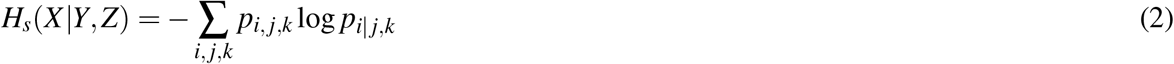

Note that, it quantifies the average residual entropy of *X* when the value of *Y* and *Z* are known.

#### Renyi Entropy

It was first discovered by Alfred Renyi^44^ in the context of information science. The Renyi entropy of the random variable *X* is defined in terms of a non-negative real number *q*, with *q* ≠ 1, as given by

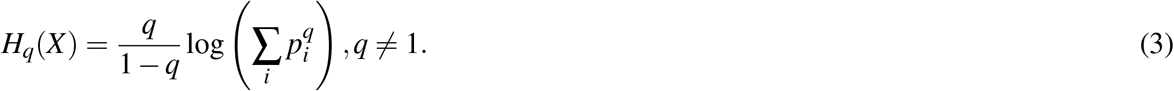

Interestingly, note that, this Renyi entropy reduces to the Shannon entropy when *q* → 1. It can also be extended for the three random variables *X*, *Y*, and *Z*, so that their joint Renyi entropy is given by

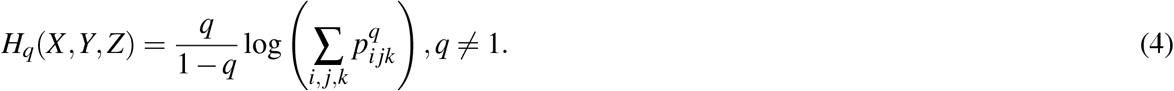

Accordingly, the conditional Renyi entropy can be defined as

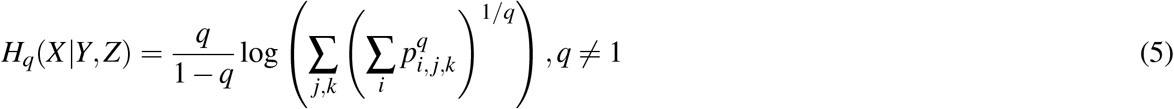

The Rényi entropy is a decreasing function of *q*. It can be showed that the conditional Renyi entropy closely correspond to a risk function, i.e. the expected error when we try to estimate the value of *X*, once we know the values of *Y* and *Z*. We refer to the associated risk as the *Renyi Risk Function* which is defined as

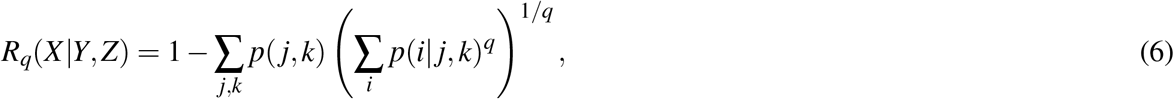

#### Renyi min-entropy

An important special case of the Renyi entropy family is the Renyi min-entropy, corresponding to the case *q* → ∞; it is also arguably the most traditional way of measuring the unpredictability of a set of outcomes. The Renyi min-entropy of *X* has the form

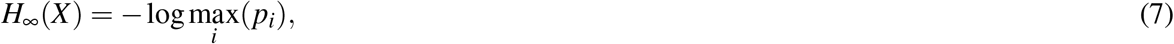

and accordingly the conditional Renyi min entropy of X given Y and Z is given by

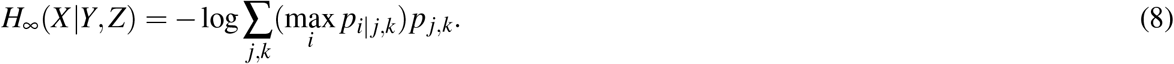

#### Tsallis entropy

It is another generalization of Shannon entropy developed from the context of statistical mechanics that yields q-normal distribution as an equilibrium probability disribution^45^.

Mathematically, the Tsallis joint entropy of the three random variables *X,Y* and *Z* is defined in terms of a tuning parameter *q* > 0 as

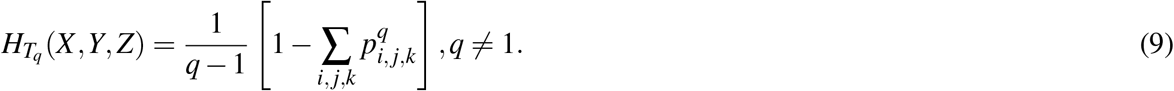

Accordingly we define the conditional Tsallis entropy of the random variable *X* given values of the random variables *Y* and *Z* as given by:

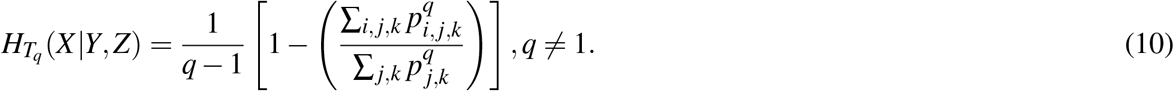

Next, we define a Tsallis risk function, i.e. the expected error when we try to estimate the value of *X*, once we know the values of *Y* and *Z*, and refer to as the *Tsallis Risk Function*. It is defined as

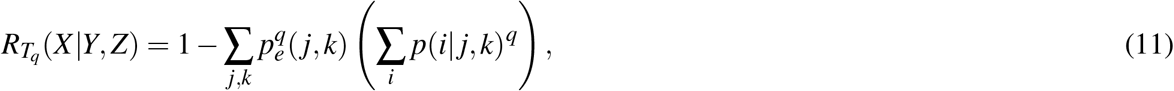

where *P*_*e*_(*j, k*) is the joint escort probability distribution^46^ given by

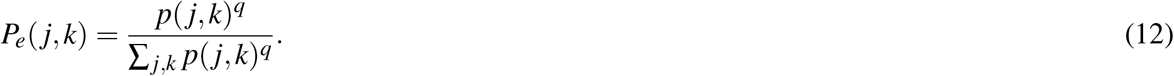

### Proposed Feature Selection Algorithm

Let, any dataset be arranged in a matrix *M*_*n*×*d*_, where *n* is number of samples and *d* is number of features. Let, *F* be the set of features, *F* = {*f*_1_, *f*_2_, *f*_3_, ..., *f*_*d*_}, and *C* be the set of classes. Our algorithm is wrapper based forward selection approach which constructs a monotonically increasing sequence {*S*} of subset of *F*. At each step, subset {*f*_*i*+1_} is acquired via the proposed algorithm according to dependency measure and added with feature subset *S* selected at the previous step. The dependency measure is evaluated using an appropriate entropy measure ℰ as described below: we will use the Shannon, Renyi and Tsallis entropy as the specific choices for ℰ.

#### Feature Relevance

Feature *f*_*i*_ is more relevant to the class label C than feature *f*_*j*_ in the context of the already selected subset *S*, when

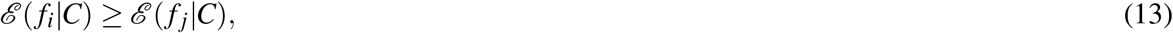

where ℰ(·, ·) is a (bivariate) conditional entropy function.

#### Feature Redundance

If feature *f*_*j*_ shares similar information with feature *f*_*i*_ than feature *f*_*j*+1_, then feature *f*_*j*_ is redundant to feature *f*_*i*_ with given information about class label *C*; it is characterized as

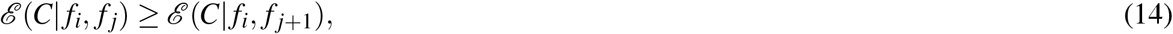

where ℰ(·|·, ·) is an appropriate conditional entropy.

#### Objective Function

We minimize the conditional entropy function between *f*_*i*_ ∈ (*F* − *S*) and *f*_*s*_ ∈ *S* (to reduce the redundancy between them) and maximize the conditional entropy function between class label *C* and *f*_*i*_ ∈ (*F* − *S*) to select the first feature, where *f*_*s*_ ∈ *S* is already selected feature.

The selected feature subset, {*S*} and the feature *f*_*i*_ ∈ (*F* − *S*) are inductively define for Renyi entropy as below:

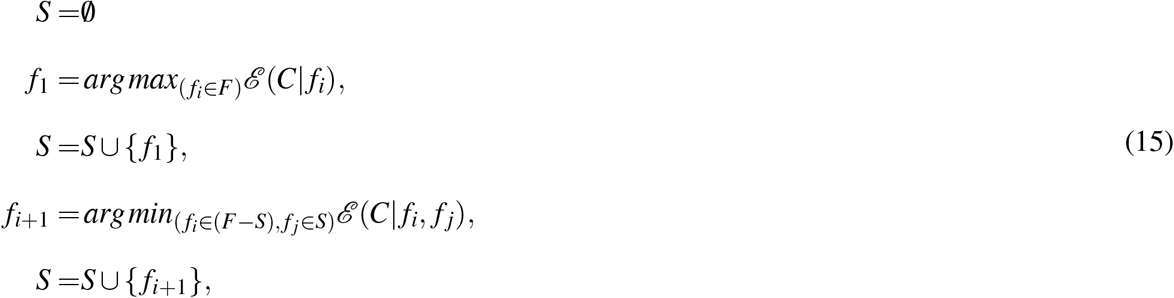

Our proposed algorithms for Renyi and Tsallis entropy are optimal in the sense of minimizing the corresponding risk functions, defined in Equation 6 and 11, respectively, these are stated by the following propositions:

##### Theorem 0.1.

*At every step, the selected feature f*_*i*+1_ *minimizes the Renyi and Tsallis risk of classification among those feature which are in selected feature subset S.*

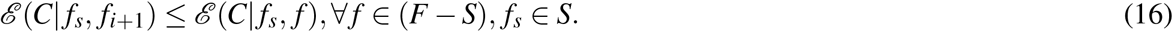

*Proof.* In order to proof Theorem 0.1, We will start from the objection function Equation 15. According to our objective function:

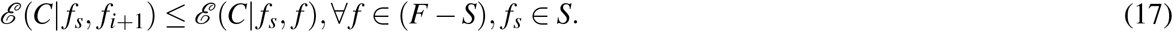

The dependency measure is evaluated using an appropriate entropy measure ℰ as described below: we will use the Renyi and Tsallis entropy as the specific choices for ℰ.

Let, *u, v′, v* represent generic value tuples and values of *f*_*s*_ ∈ *S* (Selected feature), *f*_*i*+1_ (To be selected feature at (*i* + 1)^*th*^ step), and *f* ∈ (*F* − *S*) (Non selected features) respectively. Now, after putting the generic representation in Equation 17, minimization of **Renyi risk** function will be proved.

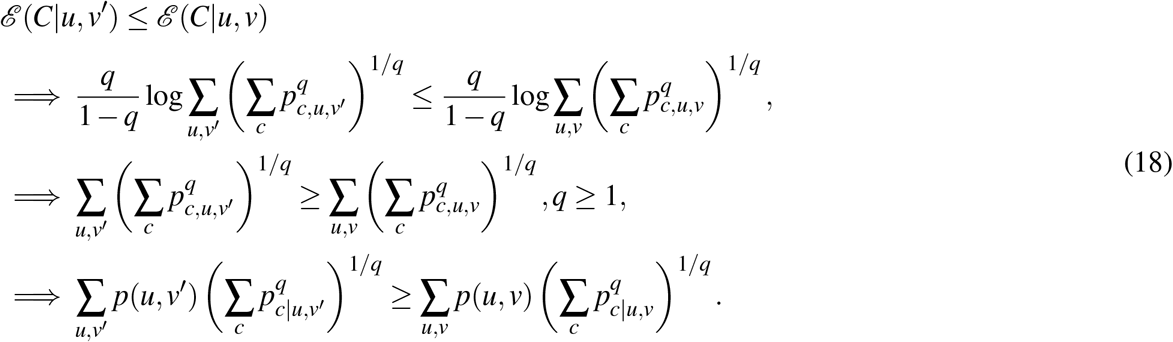

Now, for *q* ≤ 1

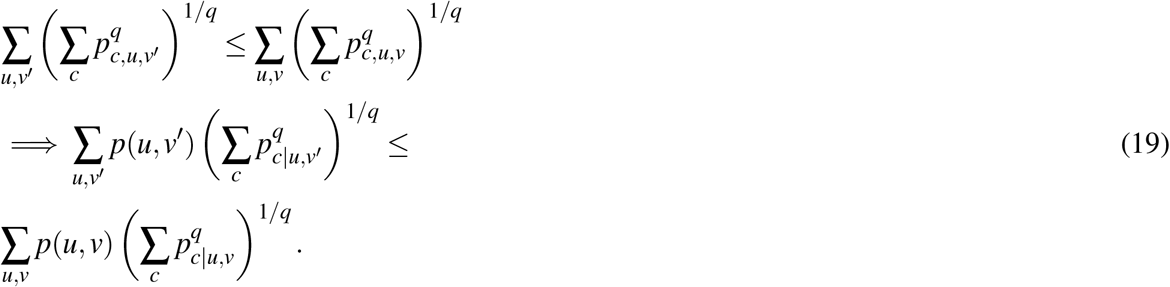

Now, Multiplying a constant 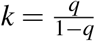 in equation 20, we get

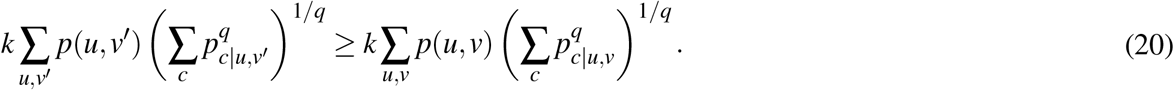

Then, by definition of Renyi risk function, described in main paper Equation 6, it can be shown that:

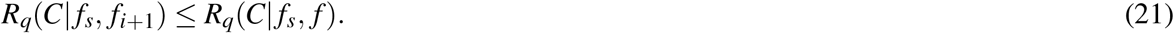

Thereafter, putting the generic representation in Equation 17, minimization of **Tsallis risk** function will be proved.

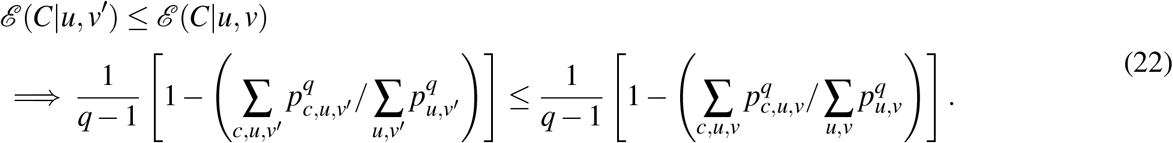

Now, for *q >* 1:

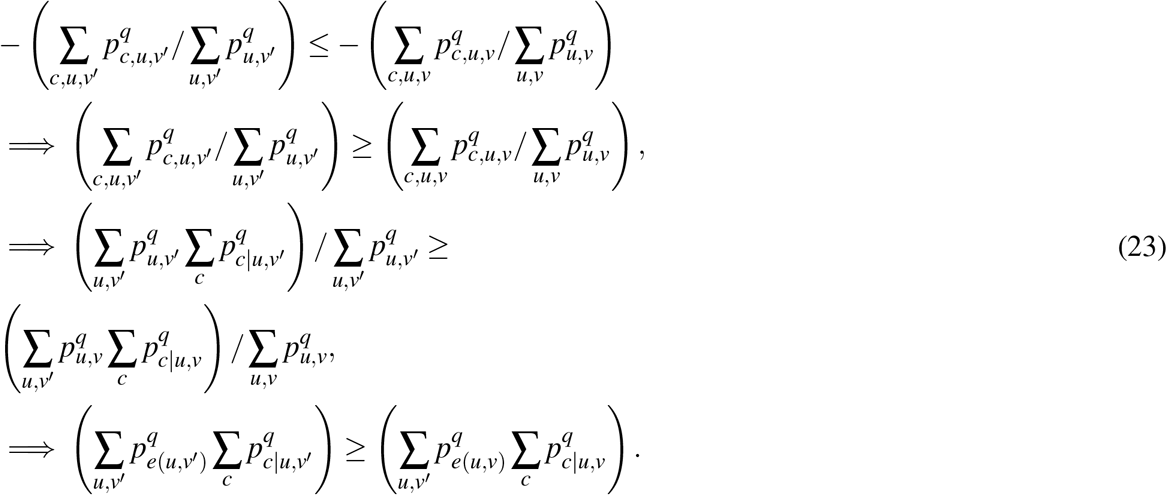

Where *p*_*e(u,v′*)_ is the Escot distribution,

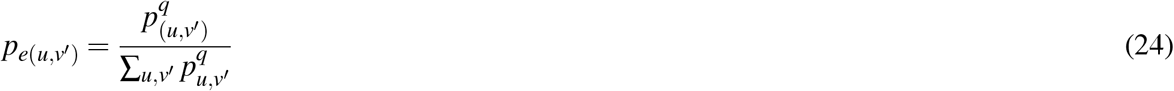

Now, for *q >* 1, the equation 22 can be written as:

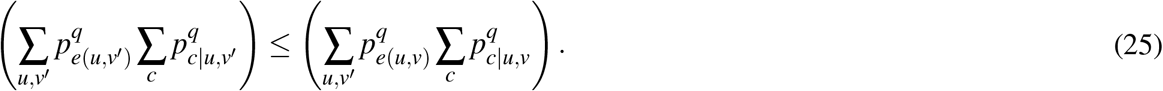

Now, Multiplying a constant 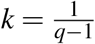 in equation 26, we get

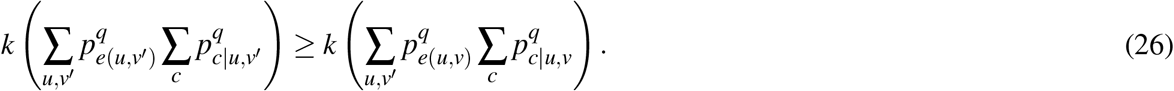

Then, by definition of Tsallis risk function, described in main paper Equation 11, it can be shown that:

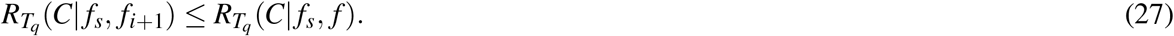

### Simulation Parameters settings in this Study

Simulation parameters for all four methods have been summarized here. ***sc-REnF*** requires the number of features (genes) to be selected as the input parameter. We select the *q*-value, 0.3 for Tsallis, and 0.7 for Renyi entropy from synthetic data simulation study, discussed in *Result* Section. The following two entropy based method are compared with the proposed method ***sc-REnF***. These are 1) Shannon entropy, 2) Min Renyi entropy. Single cell Consensus clustering (SC3)^3^ is employed to validate the informative genes. SC3 clustering is a tool for the unsupervised clustering of scRNA-seq data. SC3 achieves high accuracy and robustness by uniformly integrating different clustering solutions through a consensus approach.

### Validation Metrics used in this Study

The efficacy of the proposed method is correlated with two competitive methods on seven single cell RNA-seq datasets. To verify our proposed method (**sc-REnF**), the performance metrics are: 1) Marker Gene Selection. 2)Adjusted Rank Index (ARI) measures the similarity between two data clusters, 3) Stability using non parametric Kruskal wallis test.

## Acknowledgements

We would like to acknowledge support from JC Bose Fellowship Grant No. SB/SJ/JCB-033/201.6 dated 01/02 12017 of DST, Govt. of India.; SyMeC Project grant [BT/Med-II/NIBMG/SyMeC/2014/Vol. II] given to the Indian Statistical Institute by the Department of Biotechnology (DBT).

## Author contributions statement

SL initiated the work, SL and SR conducted the experiment(s), drafted the manuscript. AG developed the statistical theory and provided related methodological contribution. SB supervised the whole work. All authors read and approved the manuscript.

